# Quantifying the patterns of metabolic plasticity and heterogeneity along the epithelial-hybrid-mesenchymal spectrum in cancer

**DOI:** 10.1101/2021.12.18.473275

**Authors:** Srinath Muralidharan, Sarthak Sahoo, Aryamaan Saha, Sanjay Chandran, Sauma Suvra Majumdar, Herbert Levine, Mohit Kumar Jolly

## Abstract

Cancer metastasis is the leading cause of cancer-related mortality and the process of Epithelial to Mesenchymal Transition (EMT) is crucial for cancer metastasis. Either a partial or complete EMT have been reported to influence the metabolic plasticity of cancer cells in terms of switching among oxidative phosphorylation, fatty acid oxidation and glycolysis pathways. However, a comprehensive analysis of these major metabolic pathways their associations with EMT across different cancers is lacking. Here, we analyse more than 180 cancer cell datasets and show diverse associations of these metabolic pathways with the EMT status of cancer cells. Our bulk data analysis shows that EMT generally positively correlates with glycolysis but negatively with oxidative phosphorylation and fatty acid metabolism. These correlations are also consistent at the level of their molecular master regulators, namely AMPK and HIF1α. Yet, these associations are shown to not be universal. Analysis of singlecell data of EMT induction shows dynamic changes along the different axes of metabolic pathways, consistent with general trends seen in bulk samples. Together, our results reveal underlying patterns of metabolic plasticity and heterogeneity as cancer cells traverse through the epithelial-hybrid-mesenchymal spectrum of states.

## Introduction

The Epithelial to Mesenchymal transition (EMT) is a cellular programme that gives rise to a loss of epithelial phenotypes (reduced expression of cadherins involved in cell-cell attachment, loss of apical-basal polarity) with a concomitant gain in mesenchymal traits such as migration and invasion [1]. This programme has long been known to be involved in embryogenesis and wound healing in adults [2]. Cancer cells are known to activate EMT during the metastatic progression of a tumour, which allows the cells to invade and establish secondary tumours. EMT is not a binary process; instead, cells can acquire and maintain one or more hybrid epithelial/mesenchymal (E/M) states [3]. Plasticity of cancer cells along the epithelial-hybrid-mesenchymal landscape is highly dynamic, complex and multidimensional in nature [4]. Often, other relevant biological traits such as immune evasion, stemness, anoikis resistance and therapy resistance are coupled with this dynamic nature of cancer cells [5–10].

The metabolic status of cancer cells has also been reported to be coupled with EMT. Specifically, many EMT-inducing transcription factors (EMT-TFs) regulate the expression of various metabolic genes involved in glucose, lipid, glutamine, amino acid and nucleotide metabolism [11–13]. Furthermore, a change in metabolic state of cancer cells can induce a change in their EMT status [13]. The exact modalities by which these two processes are related are, however, relatively unclear, and have only begun to be investigated. For instance, activation of glycolytic enzymes by EMT has been reported in breast and prostate cancer cells [14]. A similar study reports EMT-driven activation of glycolysis in non-small cell lung cancer cells (NSCLC) by transcriptional activation of glucose transporter 3 (GLUT3) [15]. In the reverse direction, upregulation of glycolysis has been shown to promote stemness and EMT in pancreatic cancer cells [16]. On the other hand, TGF-β induced EMT has been shown to inhibit glycolysis and instead activate oxidative phosphorylation (OXPHOS) via the repression of pyruvate dehydrogenase kinase 4 (PDK4) [17], an enzyme that prevents conversion of pyruvate to acetyl-CoA. Overexpressing PDK4 inhibited EMT [17], thus demonstrating mutual regulation of these two axes of plasticity. Similarly, TGF-β induced EMT in colon cancer cells was shown to supress glycolysis by nuclear translocation of pyruvate kinase M2 (PKM2) [18], a cytosolic enzyme required for pyruvate formation. Finally, expression of SNAI1, an EMT-TF, was shown to repress another glycolytic enzyme, fructose-1,6-biphosphatase (FBP1) [19]. Similar contextdependent trends were observed in the case of the association of lipid metabolism with EMT. While TNFα or TGF-β induced EMT activation promoted the lipid synthesis pathway in prostate cancer cells [20], overexpression of SNAI1 can also lead to inactivation of lipid synthesis enzymes [21]. Overall, these studies suggest context-dependency of EMT mediated coupling to metabolic networks.

Additionally, several lipid metabolism enzymes such as acetyl-CoA synthetases (ACSL1 or ACSL4, steroyl-CoA desaturase (SCD) etc.) can activate EMT [22–24]. Notably, two pathways with whose links with EMT have been particularly well studied are glycolysis and mitochondrial metabolism. For instance, the glycolytic enzyme phosphoglucose isomerase (PGI) could also act as a cytokine and activate EMT via ZEB1 and ZEB2 stabilization in breast cancer cells [25]. But, again, this trend may not be ubiquitous across cancer subtypes and different microenvironments. The glycolytic enzyme FBP1, for instance, blocks the induction of SNAI1-driven EMT in breast cancer cells and the loss of this enzyme favours EMT, as shown *in vitro* [26]. Downregulation of several mitochondrial metabolic genes and mutations in TCA cycle enzymes have also been associated with EMT activation. Mutations in fumarate hydratase, an enzyme that converts fumarate to malate in TCA cycle, can induce EMT by inhibiting the activity of miR-200 [27]. Similarly, mutations in the TCA enzymes succinate dehydrogenase (SDH) and isocitrate dehydrogenase (IDH) also induce EMT via epigenetic suppression of miR-200, leading to alterations in the miR200-ZEB1 axis [27,28] that regulates the EMT status of cells [29]. Moreover, silencing of another TCA cycle enzyme, citrate synthase (CS), induces EMT-like cellular changes *in vitro* and promotes metastasis *in vivo* [30]. However, more recent studies reveal CS to be upregulated in several other tumour types and that its inactivation impedes EMT programme in tumour cell lines [31]. While these studies point towards a causal link between EMT and the metabolic pathways of glycolysis, fatty acid oxidation and oxidative phosphorylation, the overall interconnection landscape among these pathways is quite confounding, thereby necessitating further research.

Here, we sought to analyse the association of three main aspects of cellular metabolism – glycolysis, oxidative phosphorylation and fatty acid synthesis – with the process of EMT in more than 180 publicly available microarray/RNA-seq datasets comprising cell lines and patient tumors. We found that oxidative phosphorylation is predominantly negatively correlated with the process of EMT, and its primary regulator AMPK is primarily correlated (positively) specifically with an epithelial programme. Conversely, glycolysis and its key regulator HIF1α predominantly positively correlated with a mesenchymal programme and the induction of EMT. However, glycolysis also showed a positive correlation with the epithelial programme in many datasets, highlighting its complex interaction with the EMT programme. Fatty acid oxidation was correlated negatively with acquisition of a mesenchymal phenotype and positively with the epithelial nature of cancer cells. However, alternative modalities of association of metabolic axes with the EMT programme were also observed. Analysis of EMT induction in single cell RNA-seq data showed largely consistent trends with the generic patterns seen in the analysis of bulk samples.

## Results

### EMT scoring metrics are largely consistent across datasets

Multiple transcriptomic-based scoring metrics have been used to quantify the EMT status of biological samples [32]. We used 5 different approaches to quantify the EMT status of biological samples in a set of 182 datasets. The 76GS [33,34] and KS [35] EMT scoring methods use two different sets of gene lists (including epithelial and/or mesenchymal genes) to score a sample along the epithelial-hybrid-mesenchymal spectrum. We further used ssGSEA (single-sample Gene Set Enrichment Analysis) and/or Singscore to calculate the activity of epithelial and mesenchymal gene lists (**see Methods**) to estimate the epithelial and mesenchymal nature of the samples respectively.

For 114 out of 182 datasets, the KS score (a higher KS score implies a more mesenchymal state) is positively correlated with the enrichment of Hallmark EMT signature; in only 8 datasets, this correlation is significantly negative (**Fig 1A, left**). On the other hand, the 76GS EMT score (a higher 76GS score indicates a more epithelial state) largely correlates with epithelial signature (**Fig 1A, middle**). The KS metric was also positively correlated with an independent mesenchymal signature (**Fig 1A, right**). A comparison of 4 representative pairs of metrics show that in most datasets we looked at, these EMT scores were correlated significantly and consistently, including a negative correlation between the 76GS and KS scores, as expected (**Fig 1B**). These results show that all these EMT scoring metrics are largely consistent with one another.

**Fig 1:**
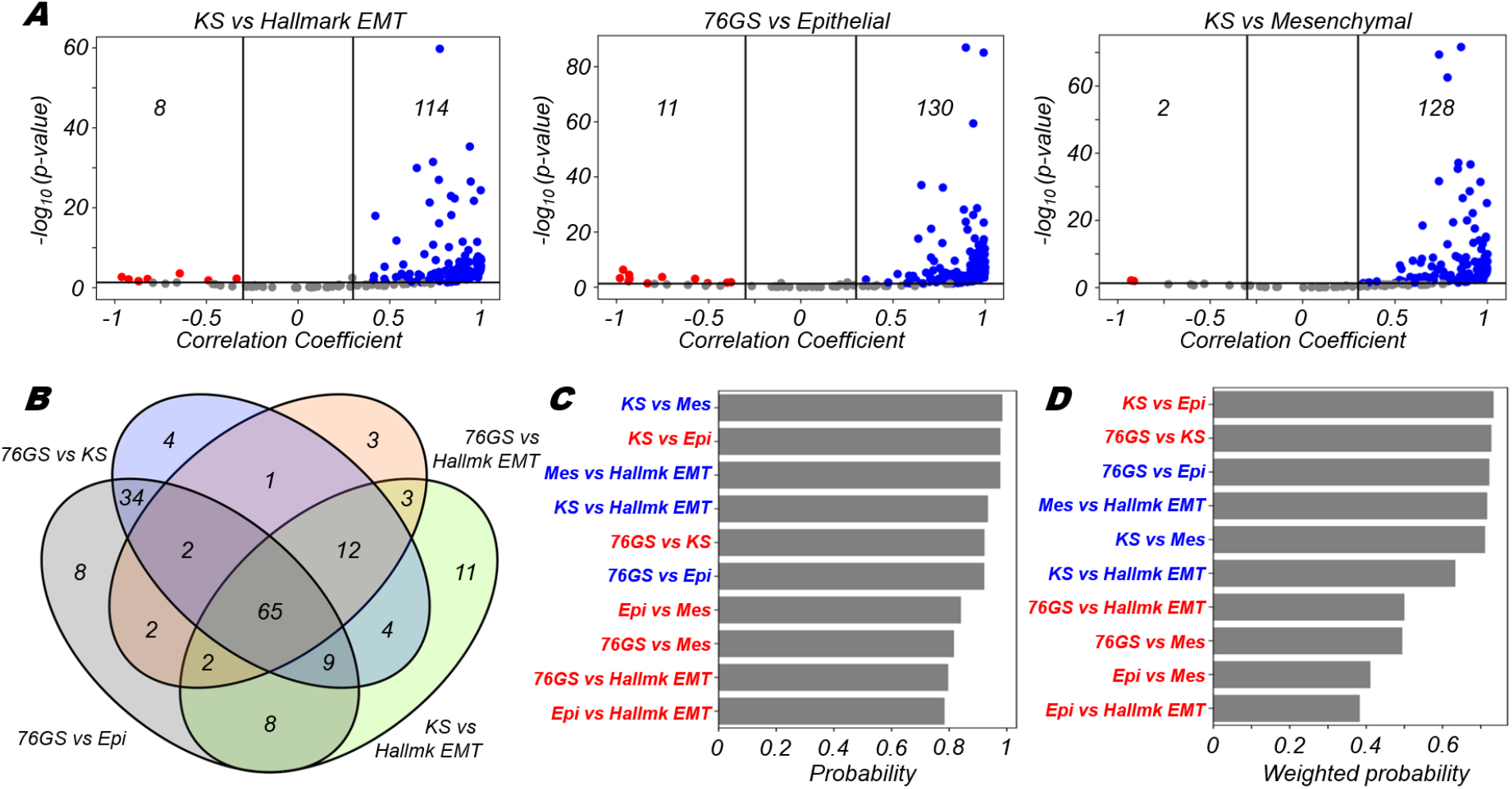
Consistency between different EMT scoring metrics. **(A)** Volcano plots depicting the Pearson correlation coefficient and the −log_10_(p-value) for 3 pairs of EMT scoring metrics – KS vs Hallmark EMT, 76GS vs epithelial and KS vs mesenchymal. Vertical boundaries are set at correlation coefficient −0.3 and 0.3. The cut-off for p-value is set at 0.05. **(B)** 4-way Venn diagram for comparison of 4 representative pairs of EMT scoring metrics. **(C)** Probability of a dataset having a positive (blue) or a negative (red) correlation (correlation coefficient > 0.3) given that it is significant (p-value < 0.05) for different pairs of EMT scoring metrics. **(D)** Probability of a dataset having a positive (blue) or a negative (red) correlation (correlation coefficient > 0.3) given that it is significant (p-value < 0.05) weighted by the fraction of significant cases for different pairs of EMT scoring metrics.

Next, we wanted to quantify the consistency of pairs of metrics if they were significantly correlated. To quantify that trend, we computed a “probability” score for a given pair of metrics by considering the number of datasets correlated significantly (p < 0.05) either positively (r > 0.3) or negatively (r < −0.3).

We computed a ratio of number of positively or negatively correlated datasets (depending on the trend seen) to the total number of datasets that showed a significant (p <0.05) association (irrespective of the direction of association). The higher this ratio is, the better the concordance between these two metrics in a given direction. We found that a) KS score vs. Mesenchymal, b) KS score vs. Epithelial and c) Mesenchymal signature vs. Hallmark EMT signatures were most consistent with one another in positive, negative and positive directions respectively (**Fig 1C**). When these probabilities were further weighted by a fraction of significant cases out of all datasets considered, we saw that a) KS score vs. Epithelial, b) 76GS score vs. KS score and c) 76GS vs. Epithelial were most consistent with each other in negative, negative and positive directions respectively, as expected (**Fig 1D**). KS score vs. Mesenchymal and KS score vs. Hallmark EMT correlations also maintained their trends as seen in earlier scenario (compare **Fig 1D** with **Fig 1C**). Put all together, these results show that the EMT metrics considered here are highly consistent with one another in a majority of the datasets.

### OXPHOS is more likely to be negatively correlated with a mesenchymal program and positively with an epithelial one

Having assessed the consistency of different EMT scoring metrics among themselves on a cohort of datasets, we next wanted to understand how the different axes of metabolism associated with EMT metrics. Oxidative phosphorylation is a predominant way by which cells generate energy for survival. To study how the biological process of oxidative phosphorylation associates with different EMT metrics, we correlated the ssGSEA activity scores calculated for the Hallmark oxidative phosphorylation gene set with different EMT metrics.

Upon correlating the OXPHOS signature with the Hallmark EMT scores, we found that although there were significant correlations both in the positive and negative directions, there were many more datasets correlated negatively (56 vs. 24) with Hallmark EMT than were correlated positively (**Fig 2A**). This overall observation is in accordance with many experimental studies that point towards a negative association between oxidative phosphorylation and EMT [36–39]. However, this relationship is not exclusive, i.e. these quantities could be positively correlated in a variety of contexts, as reported in other experimental studies [40–42]. Among all pairs of correlations between OXPHOS signature and EMT metrics, the OXPHOS-Hallmark EMT pair showed the strongest propensity of negative association with one another, given that the correlation was significant in either direction (**Fig 2B**). Furthermore, the OXPHOS-Hallmark EMT pair was also the top scoring pair when weighted with the fraction of significant cases (**Fig 2C**), further highlighting the finding that the acquisition of the mesenchymal features was more likely to result in the decline in activity of the OXPHOS gene set.

**Fig 2:**
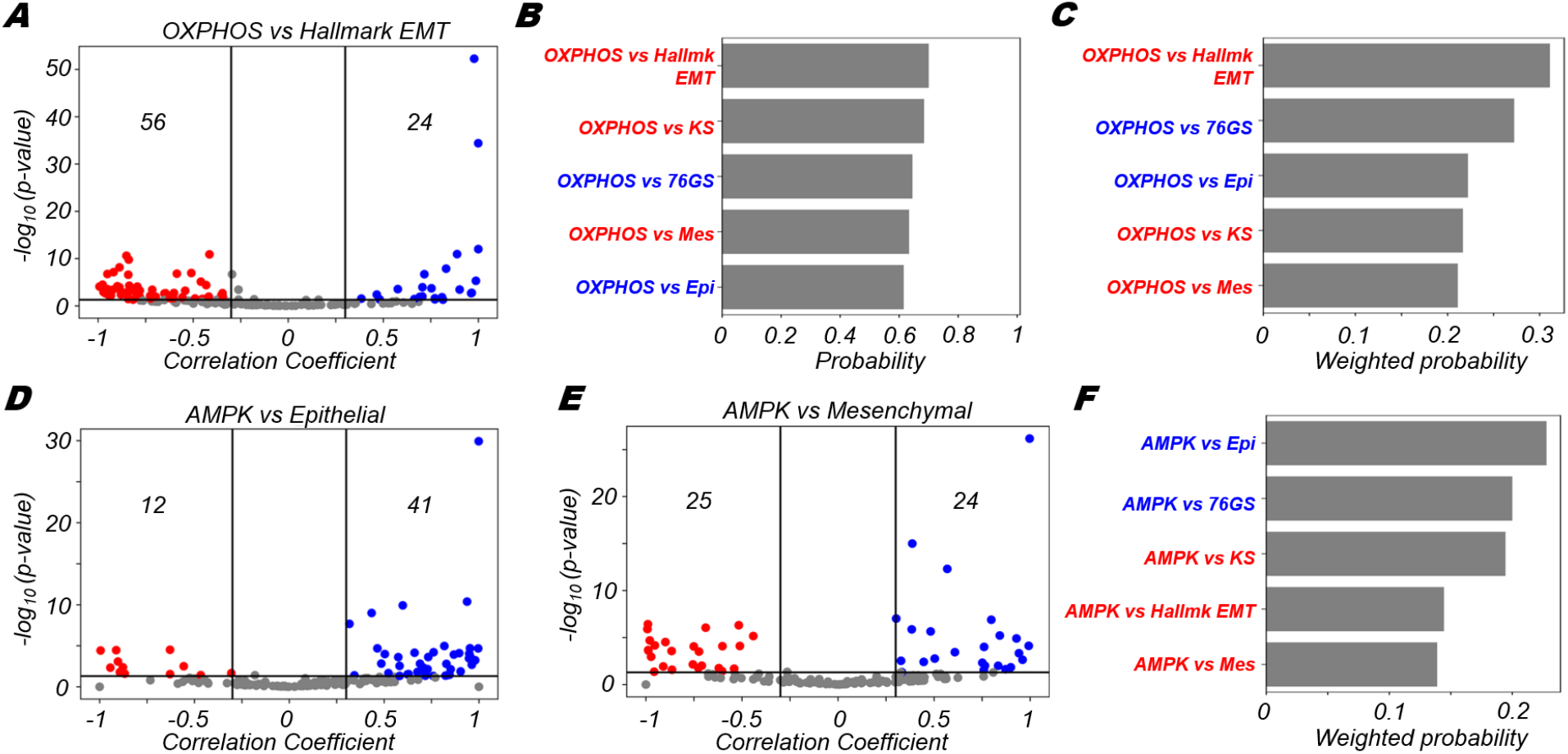
OXPHOS is more likely to correlate negatively with EMT. **(A)** Volcano plots depicting the Pearson correlation coefficient and the −log_10_(p-value) for hallmark OXPHOS and hallmark EMT signatures. **(B)** Probability of a dataset having a positive (blue) or a negative (red) correlation (correlation coefficient > 0.3) given that it is significant (p-value < 0.05) for OXPHOS and different EMT scoring metrics. **(C)** Probability of a dataset having a positive (blue) or a negative (red) correlation (correlation coefficient > 0.3) given that it is significant (p-value < 0.05) weighted by the fraction of significant cases for OXPHOS and different EMT scoring metrics. Volcano plots depicting the Pearson correlation coefficient and the −log_10_(p-value) for **(D)** AMPK signature and epithelial signature, **(E)** AMPK signature and mesenchymal signature. **(F)** Probability of a dataset having a positive (blue) or a negative (red) correlation (correlation coefficient > 0.3) given that it is significant (p-value < 0.05) weighted by the fraction of significant cases for AMPK signature and different EMT scoring metrics. Vertical boundaries for volcano plots are set at correlation coefficient −0.3 and 0.3. The cut-off for p-value is set at 0.05.

Oxidative phosphorylation in cells has been reported to be positively regulated by AMPK activity levels in cells. To assess AMPK activity, we considered a list of AMPK target genes that have been used as a proxy for the activity of the phosphorylated active form of AMPK (**see Methods**). We find that the AMPK signature is more likely to be positively correlated with epithelial signature (41 datasets showed positive correlations vs. 12 showed negative correlations) (**Fig 2D**). However, the AMPK signature did not show a strong skew towards being either positively or negatively correlated with the separate mesenchymal signature (**Fig 2E**). Together, these trends could indicate towards the fact that the active form of AMPK is likely more strongly correlated with the presence of an epithelial signature rather than with the absence of a mesenchymal one, especially if we deconvolute EMT into two-dimensional process where loss of epithelial traits and gain of mesenchymal traits can be at treated semi-independently.

Quantifying the trends of association of the AMPK signature with various EMT metrics, we noticed that the probability of positive correlation between AMPK and epithelial metrics (Epithelial signature, 76GS scores) was higher than that of a negative correlation between AMPK and mesenchymal ones (KS score, Hallmark EMT, Mesenchymal signature) (**Fig 2F**). These observations suggest that AMPK is strongly coupled with epithelial traits of cells, rather than with their mesenchymal ones. However, we noticed OXPHOS is strongly negatively correlated with both Hallmark EMT signature as well as the Mesenchymal signature (**Fig 2A, 2B**). This difference seen between trends of AMPK and OXPHOS can be due to additional context-specific factors, apart from AMPK, that might also mediate the crosstalk between EMT and OXPHOS [43], thus leading to an overall stronger negative association of OXPHOS with the hallmark EMT program.

### Glycolysis is more likely to be positively correlated with a (partial) EMT programme

Next, we wanted to check how the glycolytic process was associated with the EMT programme in the datasets we had considered. To assess this association, we correlated the enrichment (ssGSEA) scores for hallmark EMT and hallmark glycolysis signatures, across our datasets. We observed that glycolysis was more likely to be significantly positively correlated with EMT than being significantly negatively correlated (66 vs. 13 respectively) (**Fig 3A**). One would therefore expect that glycolysis should be negatively correlate with the epithelial programme or with the 76GS EMT scoring metric that assign epithelial samples a higher score. This is, however, not what we observed. Instead, glycolysis was also found to be positively correlated with scoring metrics that report an enriched epithelial program (Epithelial gene list, as well as 76GS scores), to a comparable extent with which it correlated with a mesenchymal program (**Fig 3B**). Similar trends are also seen when the association probability values were further weighted by number of significant datasets in which a given trend was observed (**Fig 3C**). Here, the association of glycolysis with Hallmark EMT programme was stronger, albeit not to a large degree, than that seen for the epithelial gene list and 76GS score, suggesting its putative association with a partial EMT state and/or other context-specific factors not included in our analysis.

**Fig 3:**
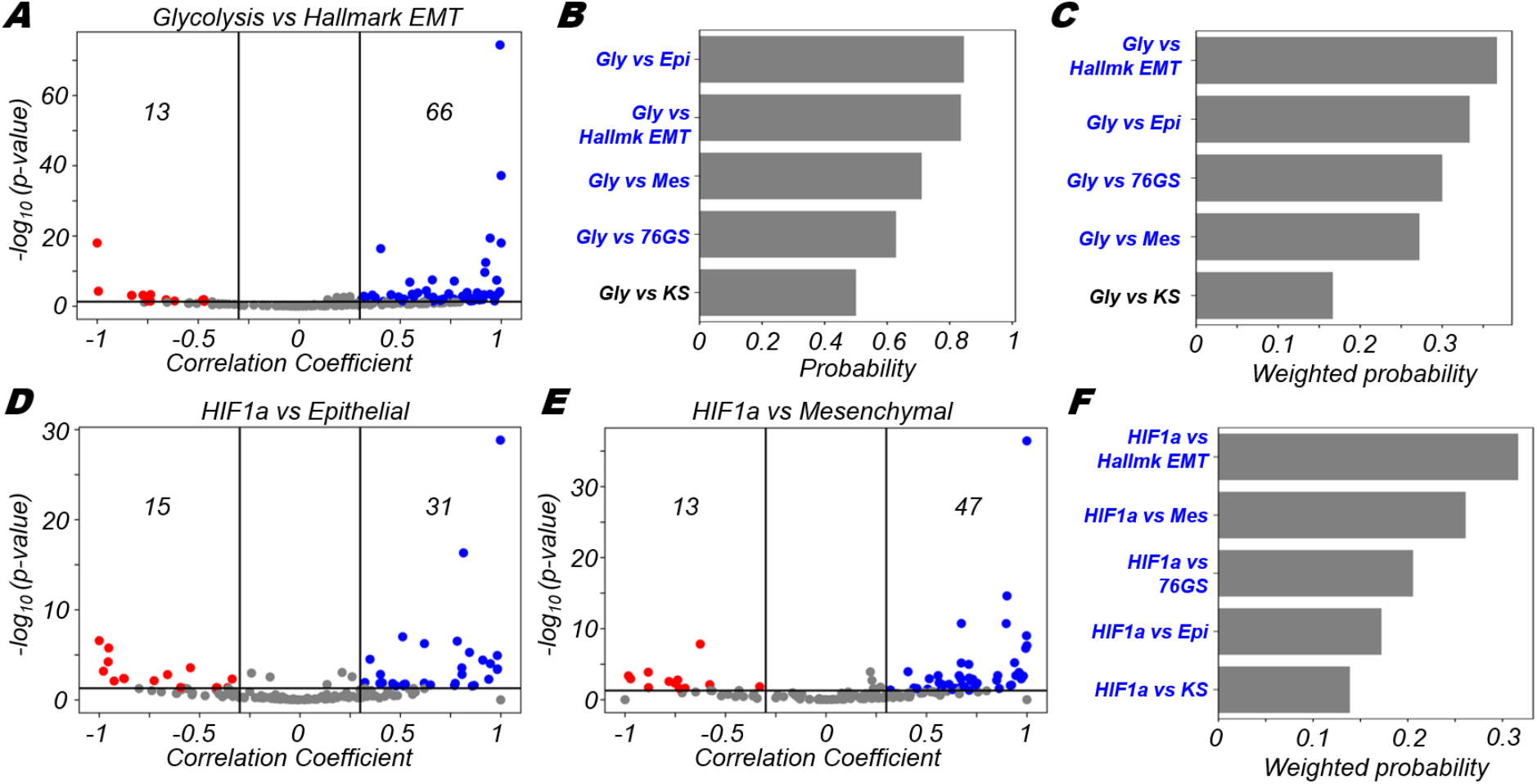
Glycolysis is more likely to correlate positively with EMT. **(A)** Volcano plots depicting the Pearson correlation coefficient and the −log_10_(p-value) for hallmark glycolysis and hallmark EMT signatures. **(B)** Probability of a dataset having a positive (blue) or a negative (red) correlation (correlation coefficient > 0.3) given that it is significant (p-value < 0.05) for glycolysis and different EMT scoring metrics. **(C)** Probability of a dataset having a positive (blue) or a negative (red) correlation (correlation coefficient > 0.3) given that it is significant (p-value < 0.05) weighted by the fraction of significant cases for glycolysis and different EMT scoring metrics. Volcano plots depicting the Pearson correlation coefficient and the −log_10_(p-value) for **(D)** HIF1a signature and epithelial signature, **(E)** HIF1a signature and mesenchymal signature. **(F)** Probability of a dataset having a positive (blue) or a negative (red) correlation (correlation coefficient > 0.3) given that it is significant (p-value < 0.05) weighted by the fraction of significant cases for HIF1a signature and different EMT scoring metrics. Vertical boundaries for volcano plots are set at correlation coefficient −0.3 and 0.3. The cut-off for p-value is set at 0.05.

HIF1α is a known mediator of the glycolytic pathway [44] and in modulating the EMT status of cells [45]. Thus, next we probed how the HIF1α signature associated with epithelial and mesenchymal programmes. Intriguingly, we found that both the volcano plots showed a skew towards the positive side (**Fig 3D-E**), suggesting that HIF1α activation may associate with a partial EMT state exhibiting both epithelial and mesenchymal features [46]. It should be noted that in the case of mesenchymal programme, the HIF1α signature was somewhat more strongly skewed towards to the positive side in comparison to the positive skew present in the case of epithelial programme (47 out of 60 datasets vs. 31 datasets out of 46 respectively) (**Fig 3D-E**). The positive association of Glycolysis as well as its known regulator HIF1α with both the epithelial and mesenchymal axes may indicate that glycolysis is a hallmark feature of hybrid E/M states. Recent observations about glycolysis accompanying collective cell migration endorse this association of glycolytic shift in partial EMT state(s) [47,48]; however, how metabolic heterogeneity maps on to leader-follower dynamic switching remains to be investigated in more detail [49–51]. Nevertheless, stronger trends as measured by weighted probability scores for HIF1α vs. Hallmark EMT and HIF1α vs. Mesenchymal compared to HIF1α vs. 76GS or HIF1α vs. Epithelial indicates enrichment of HIF1α in being associated with a relatively more mesenchymal phenotype (**Fig 3F**). The degree of coupling of gain of mesenchymal with loss of epithelial traits in a given scenario [52] may play a key role in associating HIF1α with a partial or complete EMT.

### FAO is more likely to positively correlate with an epithelial program and negatively with a mesenchymal program

Fatty acid oxidation (FAO) is a catabolic process in which fatty acids are broken down and is another key mechanism by which cancer cells can generate energy for survival [53]. Genes involved in fatty acid oxidation have been characterised previously [53]. We used one such gene set as a proxy for the activity of the FAO pathway in our datasets (**see Methods**). We found that as with OXPHOS, FAO was most likely to be negatively correlated with the Hallmark EMT programme (52 significantly negative vs. 11 significantly positive cases) (**Fig 4A**). The epithelial programme alone was more likely to be correlated positively (37 positive vs. 17 negative) (**Fig 4B**) while the mesenchymal programme alone was likely to correlate negatively (30 negative vs. 18 positive) (**Fig 4C**). These results show that FAO more likely associates negatively with acquisition of a mesenchymal phenotype. Upon calculation of the probability of positive/negative correlations given the correlation is significant as well as the overall weighted probability, we observed that the mesenchymal metrics (hallmark EMT, mesenchymal signature and KS) were all skewed towards the negative side, while the more epithelial metrics (76GS and epithelial signature) were positively correlated with FAO (**Fig 4D**). These results collectively show that while OXPHOS and FAO are more likely to be associated negatively with the mesenchymal programme and the process of EMT, glycolysis is more likely to be positively associated with the mesenchymal characteristics of cells. These associations, at least in part, are supported by the activity of molecular regulators such as AMPK and HIF1α.

**Fig 4:**
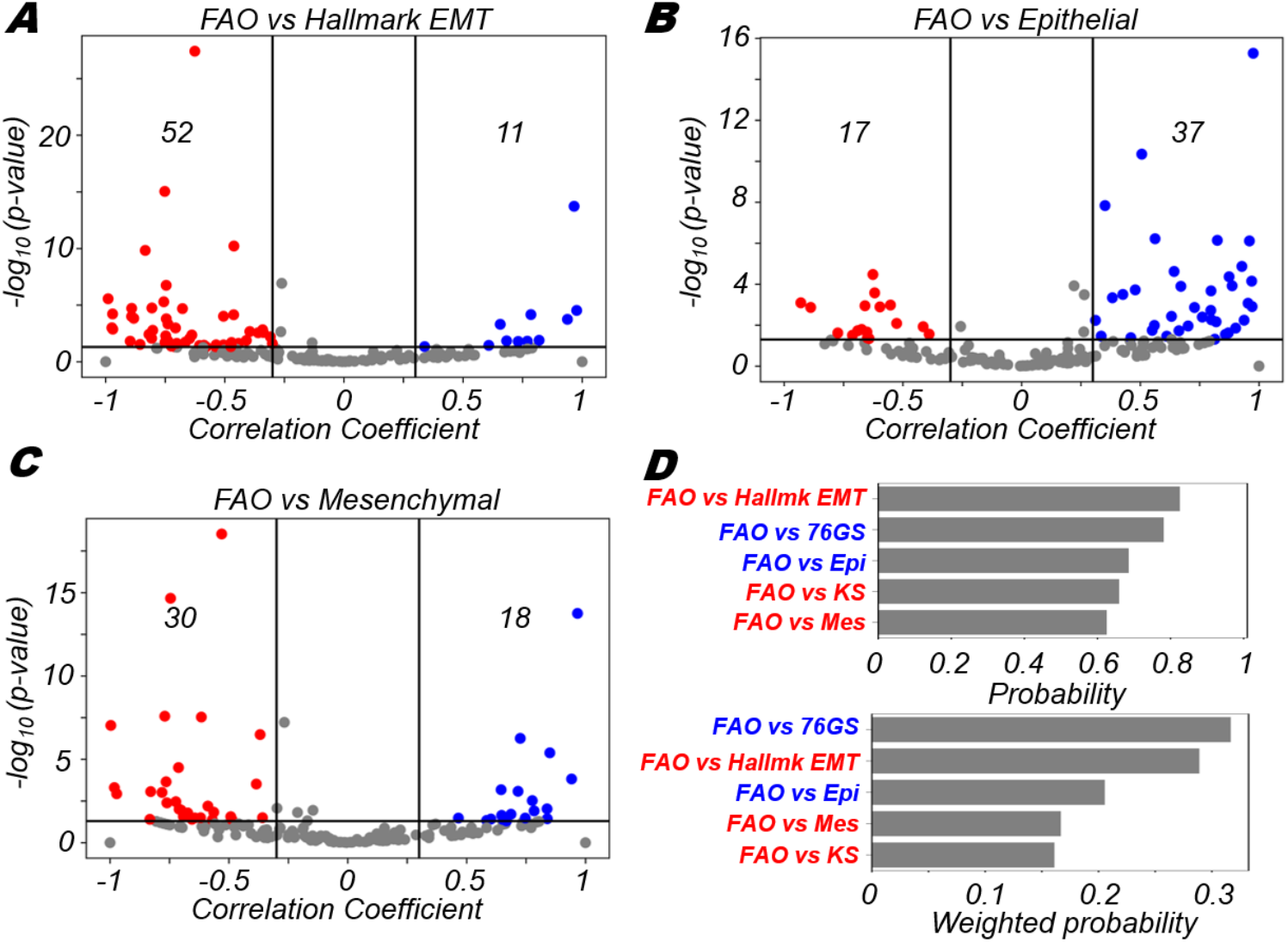
Fatty acid oxidation is more likely to correlate negatively with EMT. **(A)** Volcano plots depicting the Pearson correlation coefficient and the −log_10_(p-value) for fatty acid oxidation (FAO) and hallmark EMT signatures. **(B)** Probability of a dataset having a positive (blue) or a negative (red) correlation (correlation coefficient > 0.3) given that it is significant (p-value < 0.05) for FAO and different EMT scoring metrics. **(C)** Probability of a dataset having a positive (blue) or a negative (red) correlation (correlation coefficient > 0.3) given that it is significant (p-value < 0.05) weighted by the fraction of significant cases for FAO and different EMT scoring metrics. Volcano plots depicting the Pearson correlation coefficient and the −log_10_(p-value) for **(D)** Probability of a dataset having a positive (blue) or a negative (red) correlation (correlation coefficient > 0.3) given that it is significant (p-value < 0.05) (top panel) and weighted by the fraction of significant cases (bottom panel) for FAO signature and different EMT scoring metrics. Vertical boundaries for volcano plots are set at correlation coefficient −0.3 and 0.3. The cut-off for p-value is set at 0.05.

### Different modalities of association between pairs of metabolic pathways and EMT

After exploring the three major metabolic axes (OXPHOS, glycolysis and FAO) independently in terms of their association with EMT, we wanted to investigate the pair wise associations between the metabolic pathways in the context of EMT. For datasets under consideration, we first computed the fractions of datasets that had none, one, two or all three axes of metabolism associated significantly with the hallmark EMT signature (**Fig 5A**). Most datasets (~60%) had a maximum of one axis of metabolism correlated with the Hallmark EMT programme (**Fig 5A**). In about 25% of the datasets, hallmark EMT was not correlated with any of the metabolic axes, probably indicative of biological scenarios where these metabolic axes are not coupled directly with the EMT spectrum. In the remaining 40% of datasets, where two or more than two axes were significantly correlated with the hallmark EMT signature, we investigated if certain combinations of associations were more likely than others in context of their correlations with EMT. To answer this question, we first plotted all 45 datasets that had significant correlations with the EMT axis and either OXPHOS or glycolysis (**Fig 5B**). Among those, 21 (46.67%) datasets showed a positive correlation between glycolysis and hallmark EMT programme, while OXPHOS was negatively correlated with the hallmark EMT programme (**Fig 5B, green box**). This co-occurring association of OXPHOS and glycolysis with EMT in inverse directions has been reported earlier experimentally [54,55].

**Fig 5:**
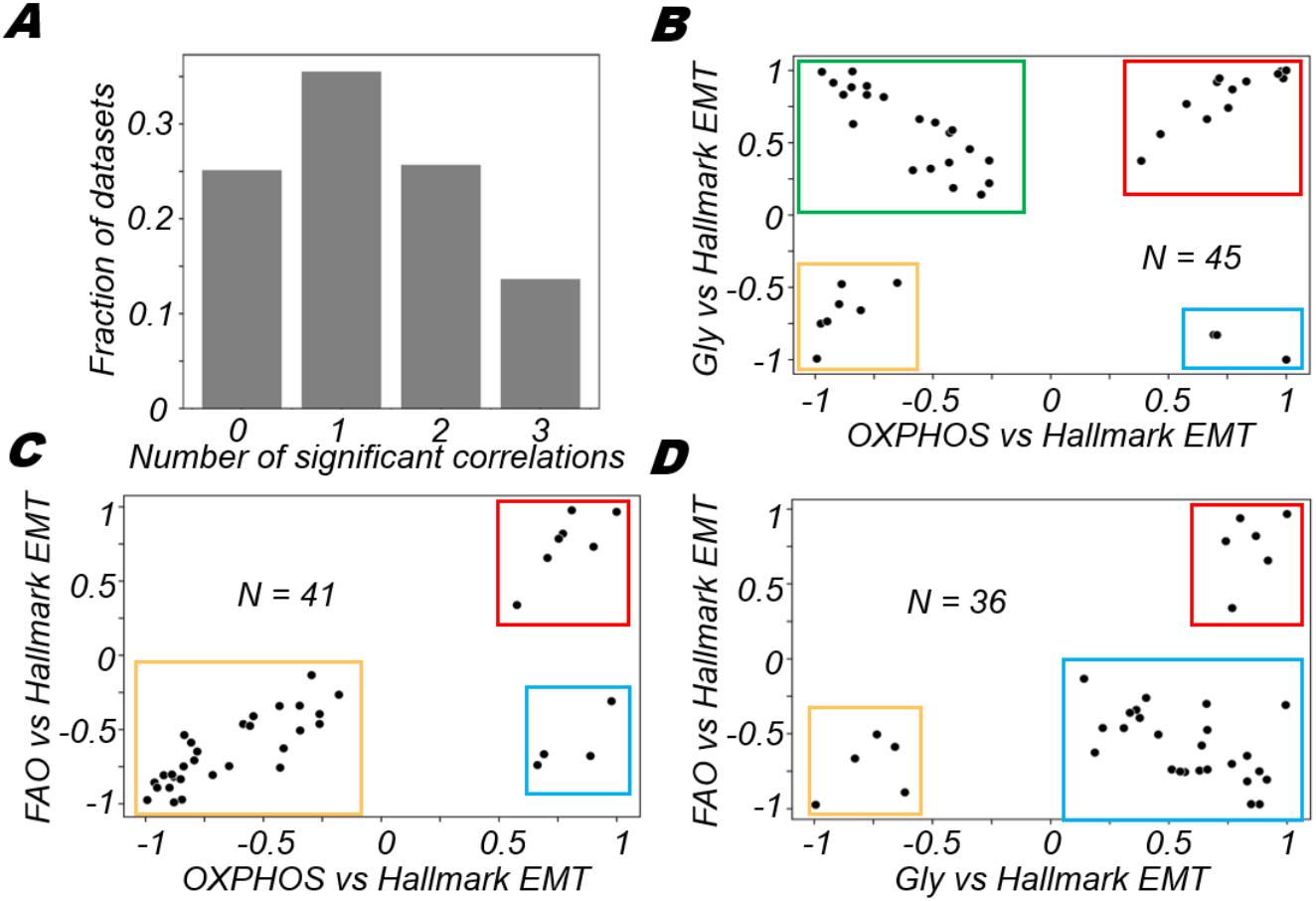
Varied associations between different metabolic axes with EMT. **(A)** Proportion of datasets that have a given number of metabolic axis significantly correlated with the Hallmark EMT signature (p-value < 0.05). Scatter plot of correlation coefficients of **(B)** OXPHOS with EMT and glycolysis with EMT **(C)** hallmark glycolysis with EMT and glycolysis with EMT **(D)** glycolysis with EMT and fatty acid oxidation with EMT.

However, this co-occurrence is not the only mode of association between these three axes. The next most predominant modality of association is the scenario where both glycolysis and OXPHOS are positively correlated with EMT – this trend is shown in 14 (31.11%) datasets (**Fig 5B, red box**). This could be indicative of the EMT associating positively with a hybrid metabolic state in which both OXPHOS and glycolysis are high [54]. The other two case – both OXPHOS and glycolysis correlating negatively with EMT (**Fig 5B, yellow box**) and OXPHOS being positively associated while glycolysis being negatively associated (**Fig 5B, blue box**) – were 15.55% and 6.67% respectively. Collectively, this analysis shows that besides predominant modalities of association between OXPHOS and glycolysis in the context of EMT, other modalities also exist although less frequently.

Next, we wanted to explore how FAO and OXPHOS are associated with each other in the context of their correlations with EMT. Upon plotting scatter plots, similar to what we had done for glycolysis and OXPHOS, we find that only 3 quadrants are populated with different propensities (**Fig 5C**). The most predominant modality was the scenario in which EMT was negatively correlated with both OXPHOS and FAO in 30 (73.17%) datasets (**Fig 5C, yellow box**). In 7 (17.07%) datasets, OXPHOS and FAO were both positively correlated with the hallmark EMT signature (**Fig 5C, red box**). In the remaining four datasets, OXPHOS was positively correlated with EMT while FAO was significantly negatively correlated with EMT (**Fig 5C, blue box**). These results show that FAO and OXPHOS are more likely to coordinated in a similar manner – either positively or negatively correlated to EMT.

Similarly, when we compared glycolysis with FAO, we found that the major modality of action was the scenario where glycolysis was positively correlated with EMT while FAO was negatively correlated with the EMT programme (**Fig 5D, blue box**). This association was observed in 25 (69.44%) datasets. The other two observed modalities of regulation were the cases where both FAO and glycolysis were both positively (**Fig 5D, red box**) or both negatively (**Fig 5D, yellow box**) correlated with EMT. Overall, our analysis uncovers the different possibilities and propensities by which these three axes of metabolism might associate with one another in terms of their connection with EMT.

### Heterogeneity in associations between different axes of metabolism in relation to EMT is also reflected in single cell RNA-seq data

Until now, our analysis was focused on bulk samples. We next examined whether there was evidence for the observed heterogeneity of association of metabolic axes with EMT present in single-cell data as well. To do so, we analysed single-cell data across different cell lines induced to undergo EMT by TGFβ [56], to ask a) if there was a shift along any of the metabolic axis upon induction of EMT, and b) whether the different modalities of associations were seen across different biological conditions. For the cell line A549, as EMT was induced for 7 days, there was a distinct rise in the hallmark EMT scores of the cells at day 7 compared to day 0 (**Fig 6A**). Concomitantly, there was a significant rise in the levels of glycolysis and FAO, while there was a drop in the levels of OXPHOS (**Fig 6A**). In the case of DU145 cells, as the hallmark EMT scores of cells increased at day 7 compared to day 0, a significant increase in the level of glycolysis, but a significant shift towards reduced levels of OXPHOS as well as FAO was noted, as expected from the most dominant trend seen in bulk datasets (**Fig 6B**). In the case of MCF7 cells, there was no significant change in glycolysis levels at day 0 and day 7. However, there was significant drop in the levels of both OXPHOS and FAO (**Fig 6C**). These results demonstrate that some level of heterogeneity in the modalities by which pairs of metabolic axes associate contingent upon the biological context, in this case the cell line considered and extent to which EMT was induced.

**Fig 6:**
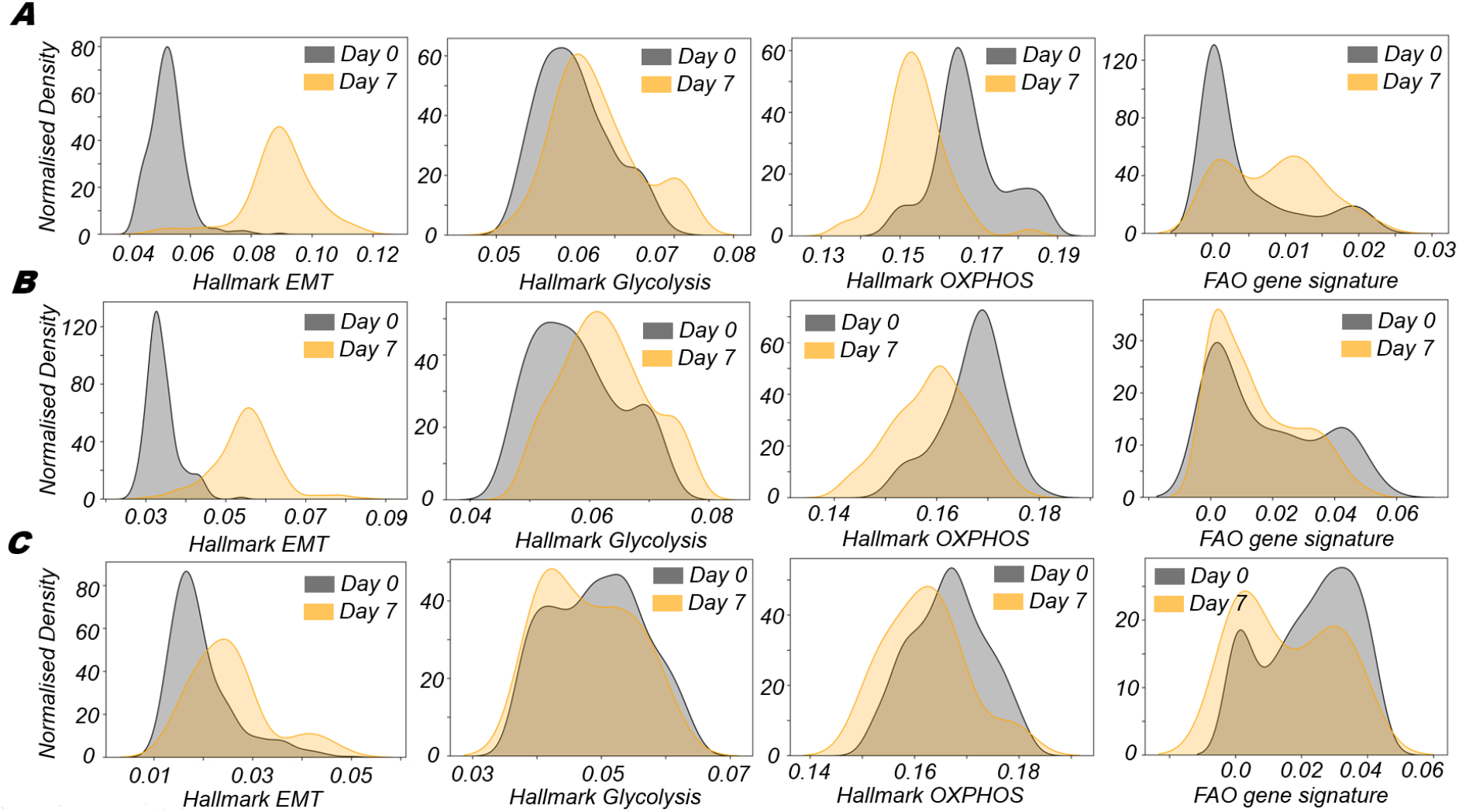
Metabolic signatures in single cells RNA-seq data upon EMT induction. Kernel density estimate plots of hallmark EMT and metabolic signatures – glycolysis, OXPHOS and fatty acid oxidation – at day 0 (untreated) and day 7 (upon TGFβ treatment) for different cell lines **(A)** A549 **(B)** DU145 **(C)** MCF7

## Discussion

Metabolic reprogramming in cancer cells is a key step in the adaptation and survival of cancer cells in the changing milieu of the tumour microenvironment. Metabolic reactions are more likely to act on a smaller time scale compared to transcriptional and translational processes. This difference in time scales makes metabolic remodelling an attractive mode for instantaneous adaptation for cancer cells. However, changes to the metabolic programmes in cells also happens at a time scale longer than such immediate adaptations. Long term changes in metabolic programmes can happen due to cross talk with other dynamic biological process in cells. In the context of EMT, cells switch from an epithelial to a more mesenchymal phenotype through multiple intermediate states, which facilitate metastasis. Changes in cellular motility and stresses arising from new microenvironments during metastasis may require more energy to adapt and survive and thus necessitate altered metabolism. Such metabolic alterations that affect the overall energy balance in the cell ultimately determine its fitness. Thus, it is not surprising that EMT and metabolism have been shown to influence one another [54,57].

In this study, we have focused on three major energy producing metabolic processes – glycolysis, oxidative phosphorylation (OXPHOS) and fatty acid oxidation (FAO). Cancer cells typically facilitate glycolysis as their primary energy source, irrespective of the presence of oxygen [58]. This process is referred to as Warburg effect or aerobic glycolysis. Conversely, the role of oxidative phosphorylation has also been observed to be important in cancer cells and cannot be ignored [59,60]. EMT induced metabolic alterations have been an active field of research with the identification of numerous mechanisms by which the metabolic state of cells is altered. Several EMT-inducing signals and EMT-TFs have been shown to activate glycolysis [15,19,61]. Further, glycolysis has been shown to promote EMT in turn, thus forming a positive feedback loop [62–65]. In addition, several studies have shown that EMT-TFs can also inhibit mitochondrial respiration and oxidative phosphorylation [66,67]. Thus, EMT has been consistently shown to be associated with activation of glycolysis and inhibition of OXPHOS, as also noted in scenarios of EMT induction [36]. However, in contrast to these studies, other pieces of evidence suggest that cancer cells with activated EMT may also have increased levels of OXPHOS in some cases [21,60]. Such conflicting findings [21,60] regarding glycolysis vs. OXPHOS may be due to differing tumor microenvironments in these studies or due to differences in cell lines/ patient samples used. Another possible explanation is that cancer cells may exhibit a hybrid metabolic phenotype [68] where both glycolysis and OXPHOS states may co-exist which allows high metabolic adaptability.

Consistent with the existing literature, our analysis of more than 180 gene expression datasets reveals that the process of EMT is more likely to be positively associated with the glycolytic process and negatively with OXPHOS programme. We also report here that FAO is also likely to be negatively associated with EMT progression, similar to OXPHOS. These broad trends are, however, not binding, but rather probabilistic in nature given different biological contexts. In other words, the absence of a universal rule indicates that cancer cells may favour glycolysis or oxidative metabolism depending on factors present in the tumor microenvironment (TME) such as availability of glucose, hypoxia, reactive oxygen species, etc. Such an ability to shift the metabolic balance dynamically may provide an advantage amidst shifting energy demands inherent in an evolving TME. Upon analysis of pairs of metabolic pathways in the context of their associations with EMT we observed that while glycolysis and OXPHOS are more likely to be antagonistic in their associations with EMT, OXPHOS and FAO were more likely to be both associated negatively with EMT. The other modalities of associations were also observed albeit in lower propensities. The observed heterogeneities were also seen in single cell RNA-seq data upon the induction of EMT.

Transcriptomic analysis of metabolic genes for given pathways, as done here, is one of the ways to estimate the level of activity of a pathway and its corresponding associations with the EMT programme. However, analysis of metabolomics data would give a more precise picture of the actual metabolic state of cells. Furthermore, there is a need to better characterise the molecular players and associated mechanisms that could allow for heterogeneity in the various modalities of associations between the different metabolic pathways and if there exist feedback loops/networks that might allow a switch from one modality of association to another. Identifying such mechanistic basis would be important to develop this understanding for any therapeutic strategies. Based on the current study, we cannot comment if the observed associations are likely to hold in the context of EMT induction only or also hold in the context of mesenchymal to epithelial transition (MET). EMT/MET dynamics has been shown to be hysteretic (non-symmetric) in nature [69–71]; whether that feature extends to metabolic reprogramming remains to be seen.

Despite these limitations, our work sheds light upon underlying patterns in terms of metabolic plasticity and heterogeneity along the epithelial-hybrid-mesenchymal spectrum in cancer cells. Understanding this coupling between EMT/MET and metabolic plasticity will enable effective targeting of cells in heterogeneous tumor populations.

## Materials and Methods

### Software and Datasets

Python (version 3.6) and R (version 4.0.2) were used for conducting all computational and statistical analyses. Microarray datasets were downloaded from NCBI GEO using *GEOquery* R Bioconductor package. FASTQ files for RNA sequencing datasets were downloaded from the ENA (European Nucleotide Archive) database. A complete list of datasets used, and the associated metadata has been provided in Supplementary Table 1.

### Pre-processing of Microarray datasets

Gene wise expression for each sample was obtained after appropriate pre-processing of microarray datasets. Probe wise expression matrices downloaded using *GEOquery* were log2 normalised and annotation files corresponding to microarray platforms were utilized for mapping the probes to respective genes. In cases where more than one probe mapped on to a single gene, the mean of expression values of all these probes was used for such genes.

### Pre-processing of RNA-seq datasets

Adapter contamination and overall quality of sequences were inspected using FastQC. Sequences were aligned with the hg38 human (or mm10 mouse) reference genome using the STAR alignment software. Finally, the raw counts for each gene were calculated with these aligned sequences using htseq-count. These raw counts were then normalised for gene length and transformed to TPM (transcripts per million) values which were then log2 normalised to obtain the final values.

### EMT scoring methods

EMT scores were calculated using five different methods for each dataset. Each method requires gene expression data as input. Each method uses a distinct gene set or a distinct algorithm.

#### 76GS

The 76-gene EMT scoring method (76GS) was developed using transcriptomic data from NSCLC cell lines and patient samples [33,34]. As the name suggests, it utilizes 76 gene signatures. The weighted sum of gene expression values of 76 genes was calculated for each sample, where the weight factors are correlation coefficients with CDH1 levels. The values obtained through this method have no specific range. EMT score for each sample was subtracted by the mean obtained from all samples such that the resultant mean score was zero. As per this new scale, negative scores indicate a M phenotype, and positive scores indicate an E phenotype.

#### KS

KS method uses the two-sample Kolmogorov Smirnov test (KS) to score EMT for cell lines and tumor samples [35]. It uses 218 gene signatures for cell line samples and 315 gene signatures for tumor samples. Briefly, cumulative distribution functions (CDFs) are obtained for each of the two signatures (E and M) and the maximum distance between these CDFs is used as the test statistic for a two-sample KS test to obtain the EMT scores. The final EMT scores lie in the range [-1, 1]. Positive & negative scores represent mesenchymal and epithelial phenotypes respectively.

#### Hallmark EMT

This method uses the hallmark geneset for EMT available (Supplementary Table 2) in the MSigDB [72] repository. For each sample, ssGSEA (single sample gene set enrichment analysis) [73] analysis was performed using this geneset to obtain the normalized enrichment score (NES). All calculations were done using the GSEAPY python library.

#### Epithelial and Mesenchymal scores

These metrics use the KS epithelial and mesenchymal gene signatures to quantify E & M status separately. A rank-based single sample scoring method called Singscore [74] was used for quantifying the enrichment level of these gene sets in a given sample. The final value obtained from this method has a range of [-1, 1]. For the epithelial score, a higher value indicates a more epithelial phenotype. The mesenchymal score also operates in a similar manner.

#### Metabolic pathways scoring methods

ssGSEA scores were calculated using the hallmark oxidative phosphorylation and glycolysis gene sets (MSigDB) to obtain OXPHOS and glycolysis signatures respectively (Supplementary Table 2). AMPK and HIF-1 signatures were quantified using expression levels of their downstream target genes as previously reported [68]. A total of 33 downstream genes for AMPK and 23 downstream genes for HIF-1 were used. The final scores were obtained using the Singscore method [74] performed on these gene sets. The FAO scores were calculated based on equations previously reported [75] which uses expression levels of 14 FAO enzyme genes.

## Acknowledgements

This work was supported by Ramanujan Fellowship (SB/S2/RJN-049/2018) awarded to MKJ by Science and Engineering Research Board (SERB), Department of Science and Technology, Government of India. HL was supported by National Science Foundation sponsored Center for Theoretical Biological Physics – award PHY-2019745, and by PHY-1605817. We are thankful to Prof. Karthik Raman (IIT Madras) for helpful discussions.

## Author contributions

MKJ and HL conceived research; MKJ supervised research; SM, SS, AS, SC and SSM performed research. SM, SS, MKJ and HL analysed data and wrote the manuscript. All authors have read and approved this version of the manuscript.

## Supplementary Table Legends

**Table S1:** Details of 184 transcriptomic datasets included in the analysis obtained from the NCBI GEO repository.

**Table S2:** List of various gene signatures used in the study for quantifying activities of the EMT program and the metabolic pathways.

## Supplementary figures

**Fig S1:**
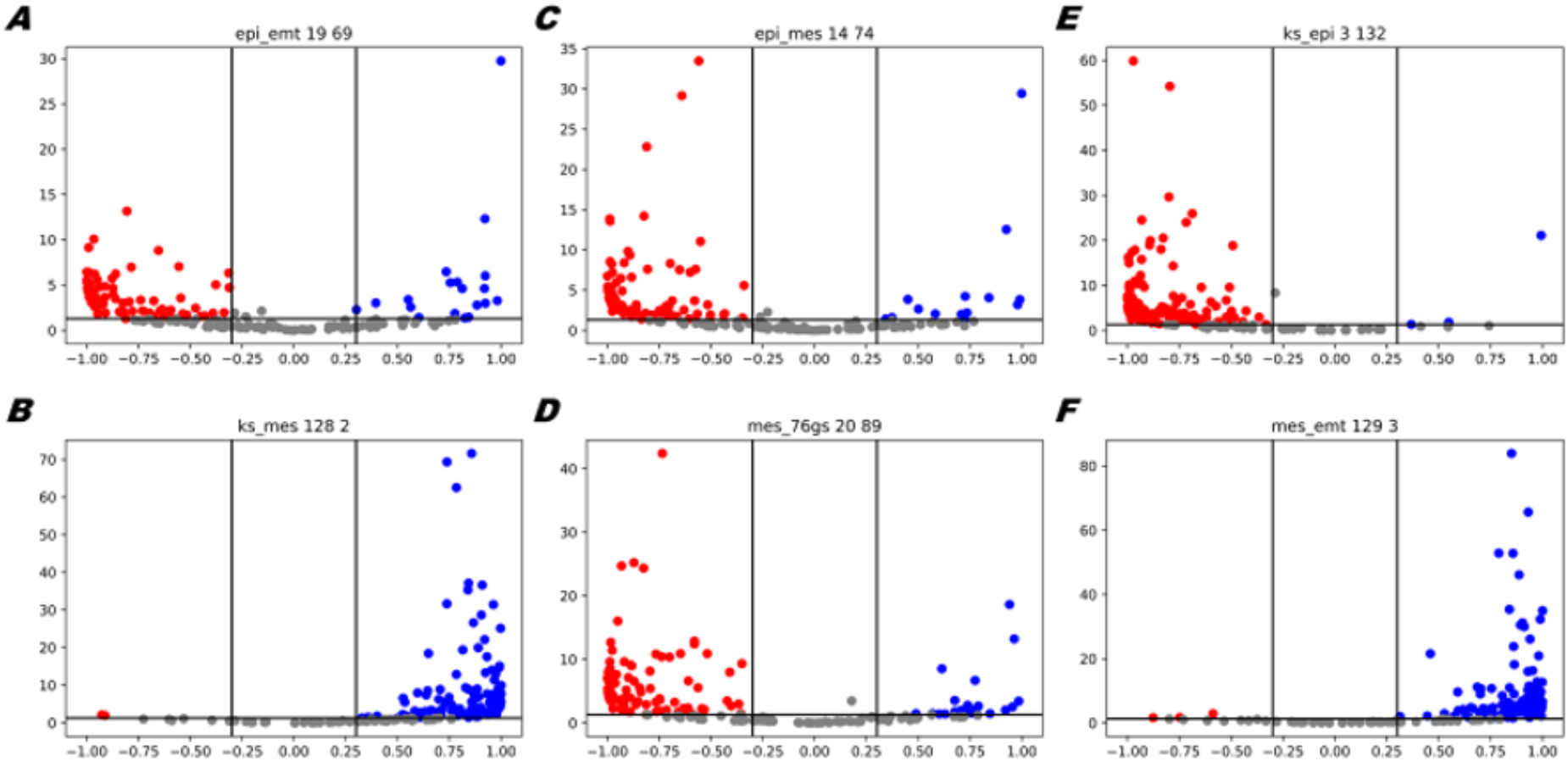
Consistency between different EMT scoring metrics. Volcano plots depicting the Pearson correlation coefficient and the −log_10_(p-value) for **(A)** Epithelial vs Hallmark EMT **(B)** KS vs Mesenchymal **(C)** Epithelial vs Mesenchymal **(D)** Mesenchymal vs 76GS **(E)** KS vs Epithelial **(F)** Mesenchymal vs Hallmark EMT. Vertical boundaries are set at correlation coefficient −0.3 and 0.3. The cut-off for p-value is set at 0.05.

**Fig S2:**
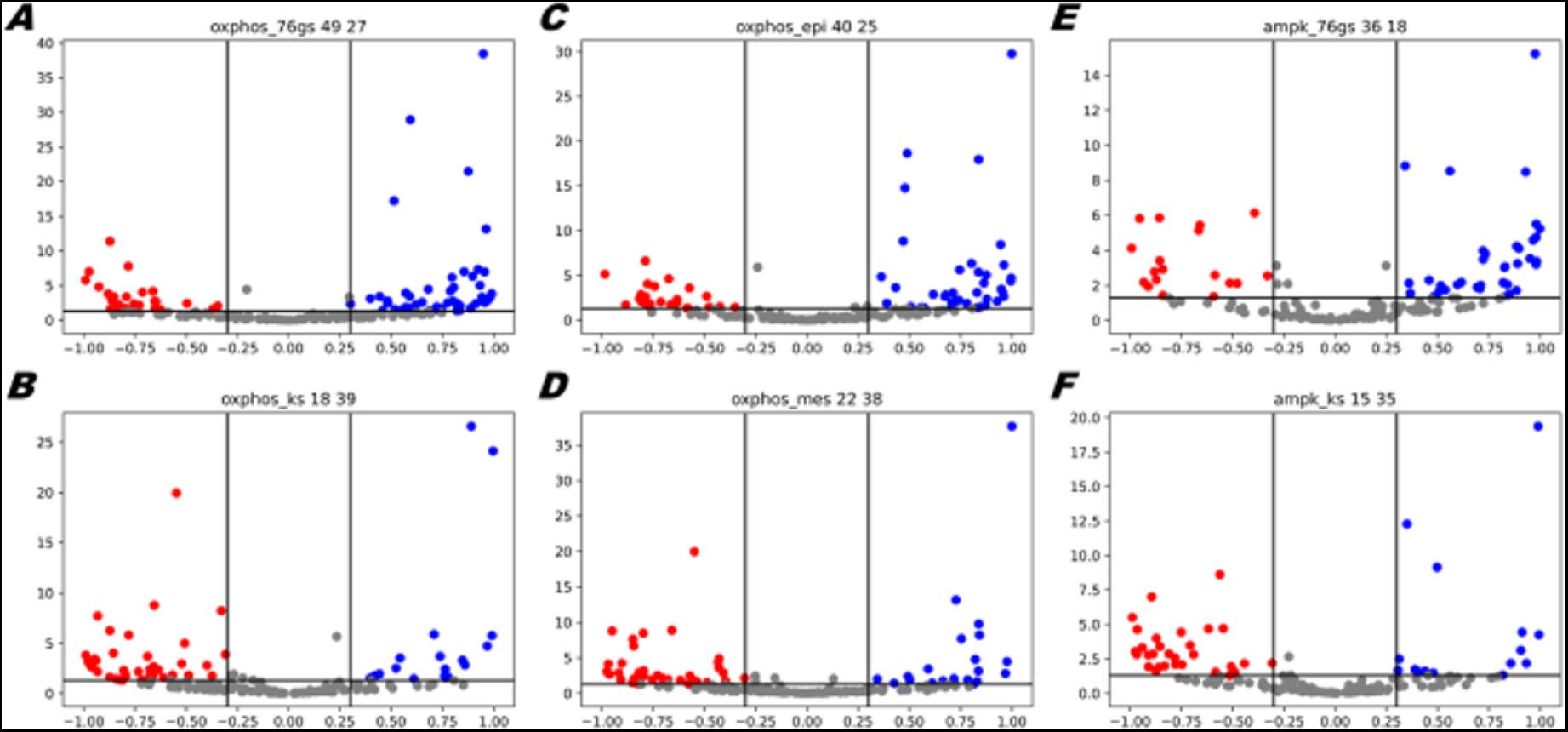
OXPHOS and its regulator AMPK is more likely to correlate negatively with EMT. Volcano plots depicting the Pearson correlation coefficient and the −log_10_(p-value) for **(A)** OXPHOS vs 76GS **(B)** OXPHOS vs KS **(C)** OXPHOS vs Epithelial **(D)** OXPHOS vs Mesenchymal **(E)** AMPK vs 76GS **(F)** AMPK vs KS. Vertical boundaries are set at correlation coefficient −0.3 and 0.3. The cut-off for p-value is set at 0.05.

**Fig S3:**
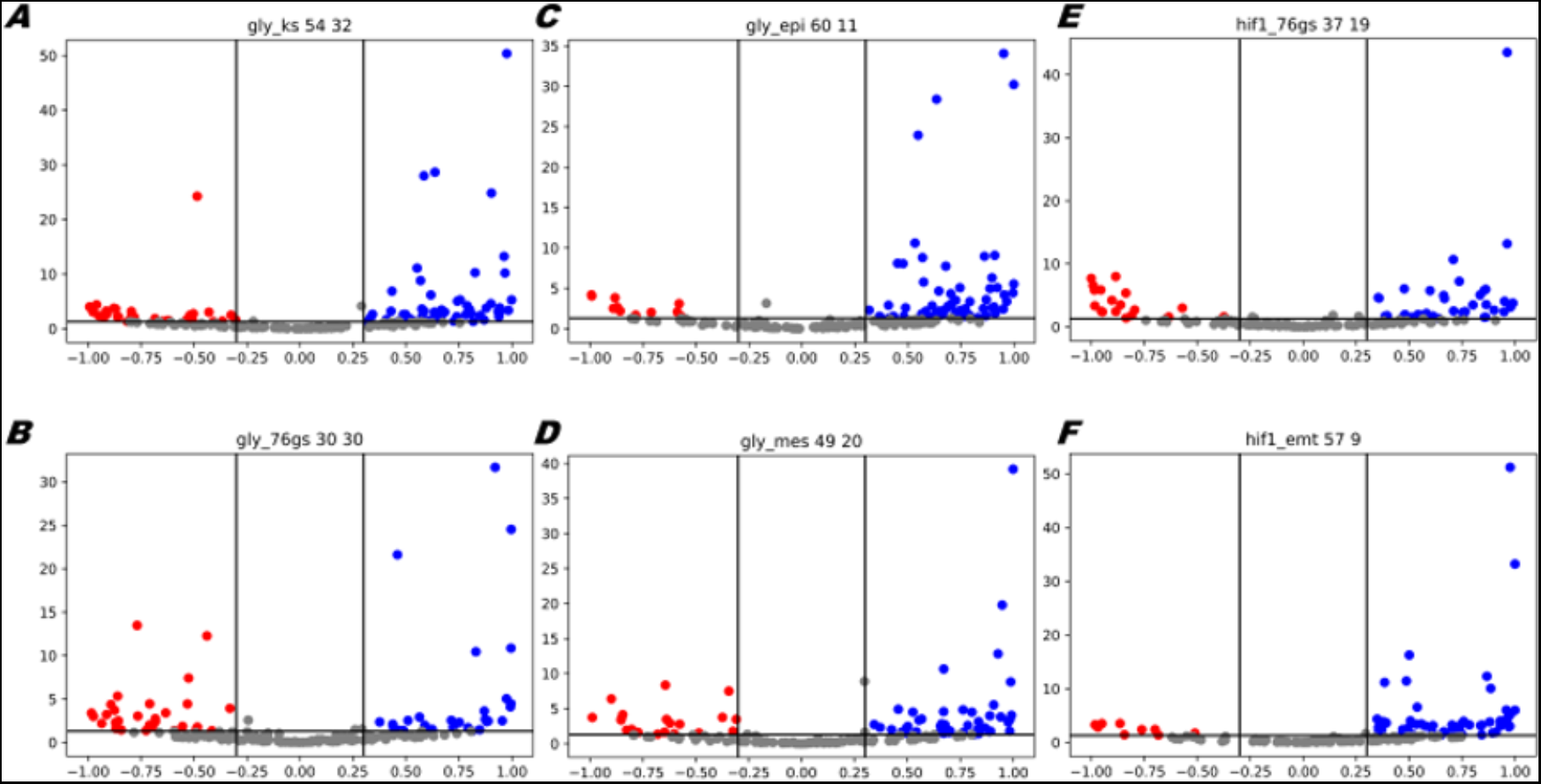
Glycolysis and its regulator HIF1a is more likely to correlate negatively with EMT. Volcano plots depicting the Pearson correlation coefficient and the −log_10_(p-value) for **(A)** Glycolysis vs KS **(B)** Glycolysis vs 76GS **(C)** Glycolysis vs Epithelial **(D)** Glycolysis vs Mesenchymal **(E)** HIF1a vs 76GS **(F)** HIF1a vs Hallmark EMT. Vertical boundaries are set at correlation coefficient −0.3 and 0.3. The cut-off for p-value is set at 0.05.

